# AutoTuner: High fidelity, robust, and rapid parameter selection for metabolomics data processing

**DOI:** 10.1101/812370

**Authors:** Craig McLean, Elizabeth B. Kujawinski

**Affiliations:** Department of Marine Chemistry and Geochemistry, Woods Hole Oceanographic Institution, Woods Hole, MA; MIT/WHOI Joint Program in Oceanography/Applied Ocean Science and Engineering, Department of Marine Chemistry and Geochemistry, Woods Hole Oceanographic Institution, Woods Hole, MA

**Keywords:** Untargeted Metabolomics, Data Processing, Data Quality, Automation, R package, BioConductor

## Abstract

Untargeted metabolomics experiments provide a snapshot of cellular metabolism, but remain challenging to interpret due to the computational complexity involved in data processing and analysis. Prior to any interpretation, raw data must be processed to remove noise and to align mass-spectral peaks across samples. This step requires selection of dataset-specific parameters, as erroneous parameters can result in noise inflation. While several algorithms exist to automate parameter selection, each depends on gradient descent optimization functions. In contrast, our new parameter optimization algorithm, AutoTuner, obtains parameter estimates from raw data in a single step as opposed to many iterations. Here, we tested the accuracy and the run time of AutoTuner in comparison to isotopologue parameter optimization (IPO), the most commonly-used parameter selection tool, and compared the resulting parameters’ influence on the quality of feature tables after processing. We performed a Monte Carlo experiment to test the robustness of AutoTuner parameter selection, and found that AutoTuner generated similar parameter estimates from random subsets of samples. We conclude that AutoTuner is a desirable alternative to existing tools, because it is scalable, highly robust, and very fast (∼100-1000X speed improvement from other algorithms going from days to minutes). AutoTuner is freely available as an R package through BioConductor.

## Introduction

Metabolomics is the study of all the compounds present within a cell, organism, or tissue. Such investigations provide a holistic snapshot of the activity within a biological matrix, and have led to a myriad of discoveries ranging from the elucidation of novel biochemical pathways, to the recognition of molecular adaptive responses to stress and the clarification of mechanisms driving cell-cell interactions.^1–3^ Advances in mass spectrometry fostered these discoveries, specifically improvements in instrument sensitivity, accuracy, and data collection capacity.^1,4,5^ Parallel advances in computational tools have historically followed to fulfill the potential of analytical improvements.^6^

Prior to data analysis, raw data from untargeted metabolomics experiments must be processed to generate a features table. Features are defined as peaks with unique mass to charge (m/z) and retention time values, with relative abundances determined by their height or area. Processing is critical to extract chemical signals from electrical noise and to correct for retention time drift across samples.^7^ A variety of untargeted data processing methods exist,^8–11^ including two commonly used tools: MZmine2^12^ and XCMS.^13^ Although these tools reliably extract true features from complex data, their performance depends on the selection of algorithmic parameters that can mitigate analytical caveats such as matrix effects and differences in analytical platforms.^14–16^ No universal set of parameters exists for all datasets, hence parameter optimization must occur prior to analysis to avoid noise inflation within the feature table.^17–19^

Tuning parameters manually is prohibitively time consuming due to the high number of possible numerical combinations. To overcome this challenge, several methods exist to identify optimal dataset-specific parameters.^20–22^ These methods each use distinct optimization functions based on maximizing or minimizing a numerical value. Each approach iteratively runs XCMS peak-picking and retention time correction algorithms until they identify a set of parameters that optimizes a desired criterion. For example, isotopologue parameter optimization (IPO), the most commonly-used parameter selection tool, scores groups of parameters by the number of features detected after XCMS that contain a naturally occurring ^13^C carbon isotopologue. Many separate XCMS runs are required to find ideal parameters, sometimes taking weeks to complete the optimization process.^20–22^ Currently, these parameter selection algorithms depend on high performance computing resources. As users continue to adopt ultra-high pressure liquid chromatography systems and rapid scanning mass spectrometers, the size and abundance of data from these platforms will inhibit the use of unscalable parameter selection algorithms by users without access to high performance computing resources.^23,24^

We designed a novel parameter optimization algorithm, AutoTuner, to ameliorate these challenges. The method performs statistical inference on raw data in a single step in order to make parameter estimates rather than iteratively checking parameter values. Further, it complements recent tools focused on generating higher-confidence feature annotations.^25–28^ AutoTuner is capable of selecting parameters for seven continuously valued parameters required for centWave peak selection algorithm used by both MZmine2 and XCMS, and it determines a key parameter for grouping in XCMS. AutoTuner is freely distributed through BioConductor as an R package.

## Theory and Design of AutoTuner

### Algorithm Overview

AutoTuner makes estimates for the following mass spectrometry peak-picking and grouping algorithms parameters: ***Group difference, ppm, S/N Threshold, Scan count, Noise, Prefilter intensity, and Minimum/Maximum Peak-width***. We chose to optimize these parameters because they have the greatest influence on the number and quality of features returned following processing and have the greatest number of possible values.^21,29^ We chose to optimize centWave peak-picking parameters over other peak-picking methods, as centWave is the recommended method for processing high-resolution untargeted data, which is increasingly becoming the standard for untargeted metabolomics.^4^ See Table 1 for a description of parameters and their matching arguments in XCMS. AutoTuner makes estimates in three steps (Figure 1):

**Table 1.** Parameters estimated through the AutoTuner algorithm. We chose to optimize these parameters due to their influence on the number and quality of features returned following XCMS data processing.^20,28^ Table S4 gives more information on these parameters.

| **Parameter** | **Description** | **XCMS parameter name** | **Functionality** | **Application** |
| --- | --- | --- | --- | --- |
| ***Group difference*** | Expected retention time deviation of an mz/rt feature between samples | bw | Grouping | XCMS |
| ***ppm*** | Parts per million (ppm) error threshold used to bin consecutive mass intensities across adjacent scans into a single peak | ppm | centWave (peak-picking) | XCMS & MZmine2 |
| ***S/N Threshold*** | The minimum ratio between peak and average noise intensity required to retain a feature | snthresh | centWave (peak-picking) | XCMS & MZmine2 |
| ***Scan count*** | Minimum number of scans required to retain a peak | Prefilter scan | centWave (peak-picking) | XCMS & MZmine2 |
| ***Noise*** | Numerical threshold used to filter out noise from true masses | noise | centWave (peak-picking) | XCMS & MZmine2 |
| ***Prefilter intensity*** | Minimum integrated intensity to required retain a peak | Prefilter intensity | centWave (peak-picking) | XCMS & MZmine2 |
| ***Peak-width*** | The width of a chromatographically resolved peak | min/max peakwidth | centWave (peak-picking) | XCMS & MZmine2 |

**Figure 1.**
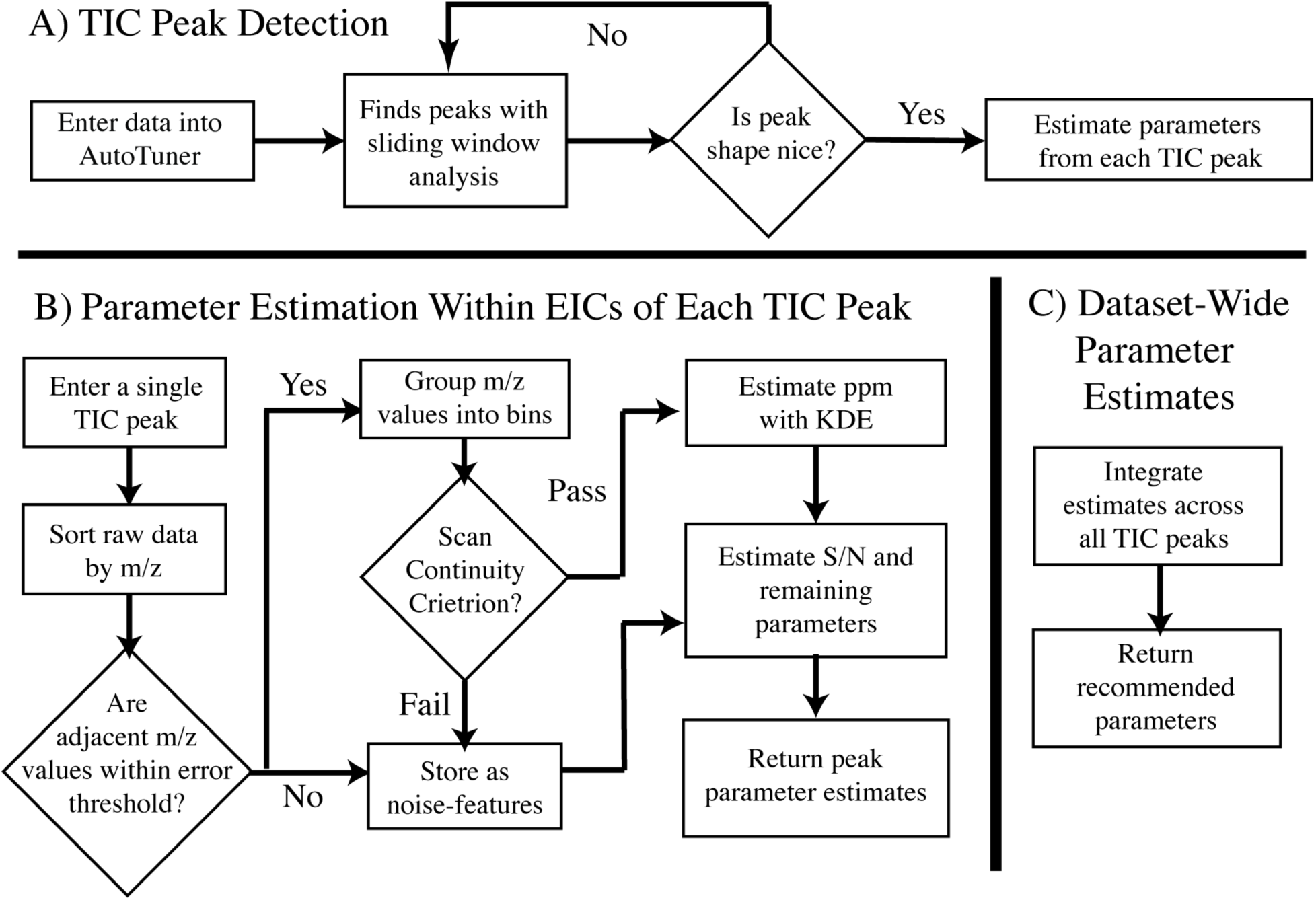
Schematic of the three stages of the AutoTuner algorithm. (A) TIC Peak Detection requires user input and is focused upon identifying peaks within each sample’s TIC. The user directly adjusts a signal processing sliding window analysis to identify peaks within the TIC. (B) Parameter Estimation Within EICs of Each TIC Peak iteratively looks at each peak to make parameter estimates from EICs. (C) Dataset-Wide Parameter Estimates aggregates results from the second stage to provide an ideal set of parameters for the entire dataset. The R package vignette at BioConductor provides an example on how to use the algorithm.

1. **TIC Peak Detection:** A user identifies peaks within each sample’s total ion chromatogram (TIC), which is the plot of integrated ion intensities entering the mass spectrometer over time.
2. **Parameter Estimation Within EICs of Each TIC Peak:** AutoTuner isolates predicted extracted ion chromatograms (EICs) within each identified TIC peak. An EIC is a plot of one or more selected m/z values in a series of mass spectra. AutoTuner applies statistical inference on all EIC peaks to estimate parameters in an unsupervised manner.
3. **Dataset-Wide Parameter Estimates:** AutoTuner integrates all peak-specific estimates into a dataset-wide set.

### Total Ion Chromatogram (TIC) Peak Detection

To identify TIC peaks, AutoTuner first applies a sliding window analysis, which detects peaks by testing if an upcoming scan’s intensity is greater than an intensity threshold determined by the average and standard deviation of a fixed number of prior scans. To ensure the correct peak bounds are retained, AutoTuner generates a linear model from the first and last three points bounding each TIC peak. If the model fails to calculate an R^2^ value greater than or equal to 0.8 or to reach a local R^2^ maximum, AutoTuner expands the ending bound by one scan and reruns the model until the model meets either criterion. The time difference of a TIC peak’s final bounds represents its width.

AutoTuner groups all TIC peaks originating from distinct samples whose maxima co-occur within each other’s retention-time bounds. It then determines the time differences between intensity maxima of all pairs of grouped peaks. AutoTuner returns the largest time difference as the estimate for the ***Group difference*** parameter that is used in the grouping step following peak-picking. Because highly complex datasets may contain distinct sample-specific peaks occurring at similar retention times, AutoTuner may overestimate this parameter. Prioritizing the inclusion of experimental replicates within AutoTuner would limit this issue. The overestimation of ***Group difference*** does not affect downstream parameter estimation, as future estimates do not involve comparisons across samples and instead focus on properties of individual EICs. At this point, AutoTuner has only collected data to estimate ***Group difference*** parameter.

### Parameter Estimation Within EICs of Each TIC Peak

AutoTuner estimates remaining parameters (***ppm, S/N Threshold, Scan count, Noise, Prefilter intensity, and Minimum/Maximum Peak-width***) from raw data contained within each individual TIC peak. A central assumption in AutoTuner is that TIC peaks represent chromatographic regions enriched in chemical ions relative to electrical noise.^7^

#### Error (ppm)

First, AutoTuner sorts all m/z values detected in mass spectra contained within the bounds of a TIC peak. AutoTuner bins m/z values if the difference in mass of two adjacent m/z values is below a user-provided threshold. AutoTuner stores unbinned m/z values as noise peaks. Because peaks of true features are made up of m/z values within adjacent scans (scan continuity criterion), AutoTuner sorts each bin by scan number to check that this criterion holds for the binned m/z values. In the case where two or more m/z values are retained from a single scan, only the m/z value with the lowest difference in mass to the previous scan’s mass is retained. If multiple m/z values occur within the first scan of the bin, the difference in mass is calculated for the next adjacent scan’s m/z value. AutoTuner stores m/z values within bins that fail the scan continuity criterion as noise peaks, similar to the noise removal step earlier.

AutoTuner estimates the parts per million (***ppm***) error parameter from the remaining bins by distinguishing between bins formed by random associations of noise peaks and those of hypothesized true features. To do this, AutoTuner first calculates the ***ppm*** of all m/z values within bins. AutoTuner then builds an empirical distribution of ***ppm*** values using a Gaussian kernel density estimator (KDE) defined by:

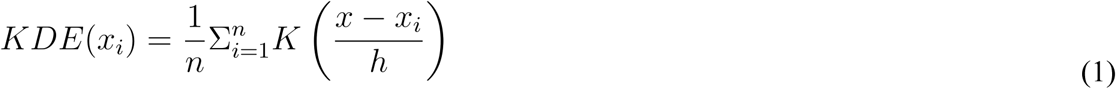

where

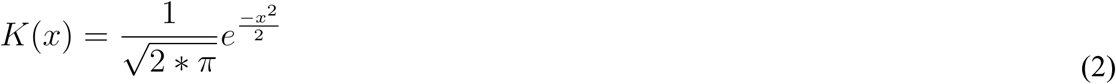

and *x* is the set of all observations, *x*_*i*_ is the ith observation, *n*is the number of observations, and *h* is a measure of smoothness for the empirical distribution. The function between ***ppm*** and absolute error is not surjective, meaning two identical absolute mass error values can have distinct ***ppm*** values. Thus, we hypothesize that the ***ppm*** value of noise peaks should be scattered widely, while ***ppm*** values of real features should be within a narrow range.^14^ Hence, we expect that by using a user-provided mass difference threshold larger than an instrument-defined threshold, the KDE will have a long-tail and a high narrow peak representing the instrument-dependent ***ppm*** of real features and a shorter smaller down-stream peak representing the ***ppm*** from erroneously binned m/z values (Figure S1).

AutoTuner subsamples the empirical distribution of all ***ppm*** values to speed downstream calculations when calculated ***ppm*** values are abundant (> 500). To do this, AutoTuner checks the similarity between the original distribution comprising all ***ppm*** values and seven distributions comprising one-half of all ***ppm*** values randomly sampled from the total. Seven was chosen arbitrarily. The distance between the original distribution and each subsampled distribution is calculated using the Kullback-Leibler divergence (KLD), a function that calculates the information theoretic gain required to describe both distributions. A KLD value of 0.5 represents an increase of one-half bit of information required to store the two distributions. If a KLD value of 0.5 or greater is not calculated across any comparison, AutoTuner replaces the original distribution with one consisting of half as many ***ppm*** values subsampled randomly from the original, and repeats the subsampling.

AutoTuner then calculates an outlier score for each ***ppm*** value to distinguish between error values derived from real features and those derived from random associations of noise using the following outlier score function:^30^

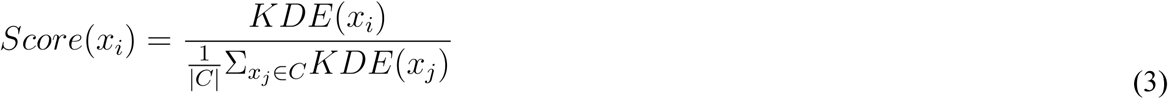

where *C* represents the largest cluster of error values and *x*_*i*_ is the ith observation similar to (1). To identify this cluster, AutoTuner uses k-means clustering, a data partitioning technique used to separate a set of observations into k-many groups. Either the gap statistic or a user-provided variance-explained threshold is used to determine the appropriate number of clusters.^31^ Using *C* ensures that the density of each calculated ***ppm*** value is normalized by the density of the true error values (Figure S1).

The ***ppm*** estimate is calculated by the following:

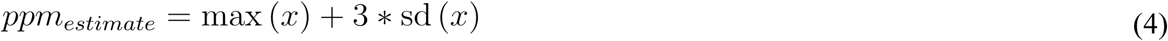

where *x* is any calculated ***ppm*** value with outlier scores above 1, and sd (*x*)is the standard deviation of all *x* values. An outlier score value above 1 indicates that the density of that particular *x* is at least as great as the expected value of the density of all elements within *C.* The statistical properties of probability distributions inspired this heuristic, as the sum of a probability distribution’s mean and three times its standard deviation provides an upper bound containing 99.7 percent of the total distribution area.^30^

#### Signal-to-Noise Threshold

We calculate the maximum intensity of each bin as well as the mean and standard deviation of the intensity of all noise features occurring within two peak widths from the original bin to estimate the signal to noise (S/N) threshold, similar to Myers et al.14 First, AutoTuner subtracts the maximum intensity of each bin (*x*_*bin,i*_) from the mean intensity of adjacent noise (*μ*_*noise*_). AutoTuner retains the bin if this difference is greater than three times the standard deviation (*σ*_*noise*_) of adjacent noise intensity values:

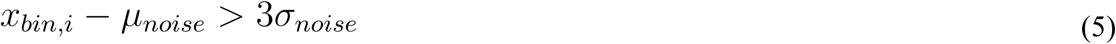

AutoTuner calculates the ***S/N Threshold*** from the smallest observed value of bin and noise intensity difference divided by the standard deviation of noise intensity across all bins passing the above threshold:

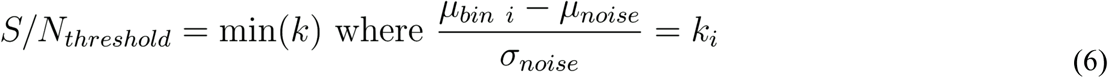

#### Remaining Parameters

AutoTuner sets the ***Scan count*** estimate as the minimum number of scans across all bins. AutoTuner estimates ***Noise*** and ***Prefilter intensity*** parameters by first determining the minimum integrated bin and single scan intensities. Then, it returns 90 percent of the magnitude of these values as the estimate to ensure that no AutoTuner-detected bin is removed during actual peak-picking. The ***Minimum Peak-width*** represents the lowest number of scans within any bin multiplied by the duty cycle of the instrument. To estimate the ***Maximum Peak-width***, AutoTuner expands bins at the boundaries of the TIC peak. The expansion continues until an adjacent scan does not contain a m/z value whose error against the mean m/z of the bin is below the estimated ***ppm*** threshold. A correlation checks to ensure that adjacent m/z values are not coming from noise after a bin has been expanded by 3 scans. For this, AutoTuner requires an absolute Spearman correlation coefficient of 0.9 between scans and intensity values for expansion to continue. AutoTuner returns the ***Maximum Peak-width*** across bins.

### Dataset-Wide Parameter Estimates

AutoTuner uses the average of all ***ppm*** and ***S/N Threshold*** values weighed by the number of bins within each TIC peak to return dataset-wide estimates for these parameters. For dataset-wide values of ***Scan count, Noise, Prefilter intensity***, and ***Minimum Peak-width***, AutoTuner returns the minimum values from all bins detected. The maximum calculated ***Group difference*** parameter represents the dataset wide parameter estimate. The average of each sample’s maximal peak-width represents the ***Maximum Peak-width*** estimate.

## Experimental Demonstration

### Materials

We chose a suite of 85 metabolites that represent compounds expected in metabolomic experiments, including cofactors, amino acids, and secondary metabolites. Of these, 41 ionized exclusively in negative mode, 28 ionized exclusively in positive mode, and 16 ionized in both modes. See Table S1 for a complete list of standards.

We prepared stock solutions of each metabolite standard in water or a 1:1 mix of methanol and water at 1000 mg mL^-1^, unless constrained by solubility. Some standards required the addition of ammonium hydroxide or formic acid for dissolution. We stored stock solutions in the dark at -20°C. We created a standard metabolite mix (10 mg ml^-1^) from the stock solutions and diluted with Milli-Q water to obtain four solutions where the concentration for each standard was 500 ng mL^-1^. We obtained standards at the highest grade available through Sigma Aldrich (MO, USA) with the exception of dimethylsulfoniopropionate (DMSP), purchased from 21 Research Plus Inc. (NJ, USA).

### Mass Spectrometry

We analyzed four replicates of the standard mixes with ultra-high-performance liquid chromatography (UPLC; Accela Open Autosampler and Accela 1250 Pump (Thermo Scientific)), coupled via heated electrospray ionization (H-ESI) to an ultrahigh resolution tribrid mass spectrometer (Orbitrap Fusion Lumos (Thermo Scientific)). We performed chromatographic separation with a Waters Acquity HSS T3 column (2.1 × 100 mm, 1.8 μm) equipped with a Vanguard pre-column, both maintained at 40°C. We used mobile phases of (A) 0.1% formic acid in water and (B) 0.1% formic acid in acetonitrile at a flow rate of 0.5 mL min^-1^ to elute the column. The gradient started at 1% B for 1 min, ramped to 15% B from 1-3 min, ramped to 50% from 3-6 min, ramped to 95% B from 6-9 min, held until 10 min, ramped to 1% from 10-10.2 min, and finally held at 1% B (total gradient time 12 min). We made separate positive and negative ion mode autosampler injections of 5 μL. We set electrospray voltage to 3600 V (positive) and 2600 V (negative), and source gases to 55 (sheath) and 20 (auxiliary). We set the heated capillary temperature to 375°C and the vaporizer temperature to 400°C. We acquired full scan MS data in the Orbitrap analyzer (mass resolution 120,000 FWHM at m/z 200), with an automatic gain control (AGC) target of 4e5, a maximum injection time of 50 sec, and a scan range of 100-1000 m/z. We set the AGC target value for fragmentation spectra at 5e4, and the intensity threshold at 2e4. We collected all data in profile mode.

### Validation Data

We used two published datasets to validate AutoTuner’s performance on experimental data: (1) a bacterial culture experiment^32^, MetaboLights^33^ identifier MTBLS157, and (2) a rat fecal microbiome, by direct contact with authors (Table 2).^34^

**Table 2.** Information on the datasets used to test AutoTuner’s performance. The mass spectrometers and liquid chromatography systems herein are some of the most commonly used analytical platforms for untargeted metabolomics.^4^

| Dataset | Reference | Access | Mass Spectrometer | Liquid Chromatography | Sample Number | Ionization Mode |
| --- | --- | --- | --- | --- | --- | --- |
| Standards | (current project) | (current project) | Orbitrap Fusion Lumos (Thermo) | Ultra-high Performance Liquid Chromatography (Accela 2015 Pump - Thermo) | 4 | Pos/Neg |
| Culture | 31 | MetaboLights MTBLS157 | Hybrid Linear Ion Trap 7T Fourier Transform Ion Cyclotron Resonance (Thermo) | High Performance Liquid Chromatography (Surveyor MS Pump Plus - Thermo) | 45 | Pos/Neg |
| Community | 33 | Contributing Author | Time-Of-Flight Tandem Mass Spectrometer (Xevo-G2 waters) | Ultra-high Performance Liquid Chromatography (Acquity - Waters) | 90 | Pos/Neg |

#### Data Processing and Quality Control

We converted all raw data files from their proprietary formats to mzML files using msconvert.^35^ All computing of mzML files took place within an Ubuntu Xenial 16.04 Google Cloud instance with 8 CPUs and 10Gb of memory. During time comparisons, we used 8 and 1 CPU(s) to obtain IPO and AutoTuner data processing parameters, respectively. We used an m/z mass error threshold of 0.005 Daltons for AutoTuner, because this absolute error was sufficiently large to return a broad range of error values (in ppm) greater than those of the mass analyzers used to generate the validation data (Table 2).

We used XCMS and centWave to generate feature tables for each dataset,^13,36^ and CAMERA for isotopologue and adduct detection.^37^ Table S2 contains parameters used for processing. For the standards, we searched for the most abundant parent ion within EICs. We confirmed the presence of a metabolite standard if a feature had an intensity above 1e4, was within an exact mass error of 5 ppm of the parent ion, and had a retention time error of 5 seconds from the EIC peak. For the culture experiment, the data was subjected to quality control as described previously.^38^ Briefly, we removed features detected in process blanks, features detected within only one replicate, and features representing isotopologues and adducts. Additionally, we removed features with coefficient of variation values above 0.4 across six pooled samples. We defined overlapping features in AutoTuner- and IPO-parameterized feature tables to be those with ppm error below 5 and retention time differences less than 20 seconds. The culture experiment allowed a higher retention time correction because it relied on data collected with HPLC compared to the standards which were analyzed with a UPLC system.

### Statistical Analyses

We applied several distinct statistical methods to summarize the various pieces of data used to validate AutoTuner. We used R programming language to perform all analyses (CRAN R-Project). We used a Kolmogorov-Smirnov Test (KS-test) to compare empirical cumulative distribution functions. We used the hypergeometric test to compare MS^2^ enrichment of IPO- and AutoTuner-specific features against features observed in the intersect of the two datasets. In order to estimate the robustness of AutoTuner parameter estimation, we performed a Monte Carlo experiment by running AutoTuner on distinct subsets of the data. We first randomly selected 7 subsets of 11 samples to compare the variability across parameters. We used the coefficient of variation from estimates within each group as a measure of variability. We also performed linear regressions on these values to find trends between estimate variability and sample numbers. We randomly selected 3 to 9 samples from each of these subsets 55 times. In total, there were 385 estimates for each group of 3 to 9 samples, resulting in a total of 2695 separate runs of AutoTuner per dataset. We performed a sensitivity analysis to determine how distinct values of mzDiff impact downstream data processing, the only continuous valued centWave parameter not optimized by AutoTuner To accomplish this, we counted the number of unique features between pairs of feature tables generated with mzDiff parameters varying by a value of 0.001.

## Results

### AutoTuner Accuracy and Comparison to IPO

At this time, the only publicly-available method for automated selection of peak-picking parameters for XCMS is isotopologue parameter optimization (IPO).^22^ IPO uses a gradient descent algorithm that requires users to iteratively run centWave with different combinations of parameters until the set that maximizes a scoring function is identified. We used 5 distinct metrics to compare the accuracy, ease of use, and downstream data quality of IPO- and AutoTuner-derived parameters. The metrics include the accuracy, number of features, the peak areas and shapes of EIC peaks detected using parameters from only one of the two methods, and MS^2^ count.

We searched for 85 known chemical standards (a total of 101 possible ions) within feature tables generated with IPO- and AutoTuner-derived parameters to test the influence of each parameter selection method on data processing accuracy (Table 2, Table S1). We detected 82 and 100 standards within the feature table generated with IPO- and AutoTuner-derived parameters, respectively. Figure S2 provides an example of compounds that were only detected with AutoTuner and were absent when the IPO-derived parameters were used.

We first compared the number of features from culture data generated with parameters from each algorithm to understand the influence of parameter selection on downstream data quality (Figure 2A, Figure S4A). Each feature table contained a distinct number of total features following processing and quality control (Table S3). In positive ion mode, AutoTuner-derived parameters detected fewer unique features (203) compared to 2606 unique features detected with IPO-derived parameters (Figure 2A), while sharing 1022 features between them. A similar situation was observed in negative ion mode where AutoTuner detected 540 unique features compared to 3420 unique features found with IPO-derived parameters, while sharing 904 features (Figure S4A).

**Figure 2.**
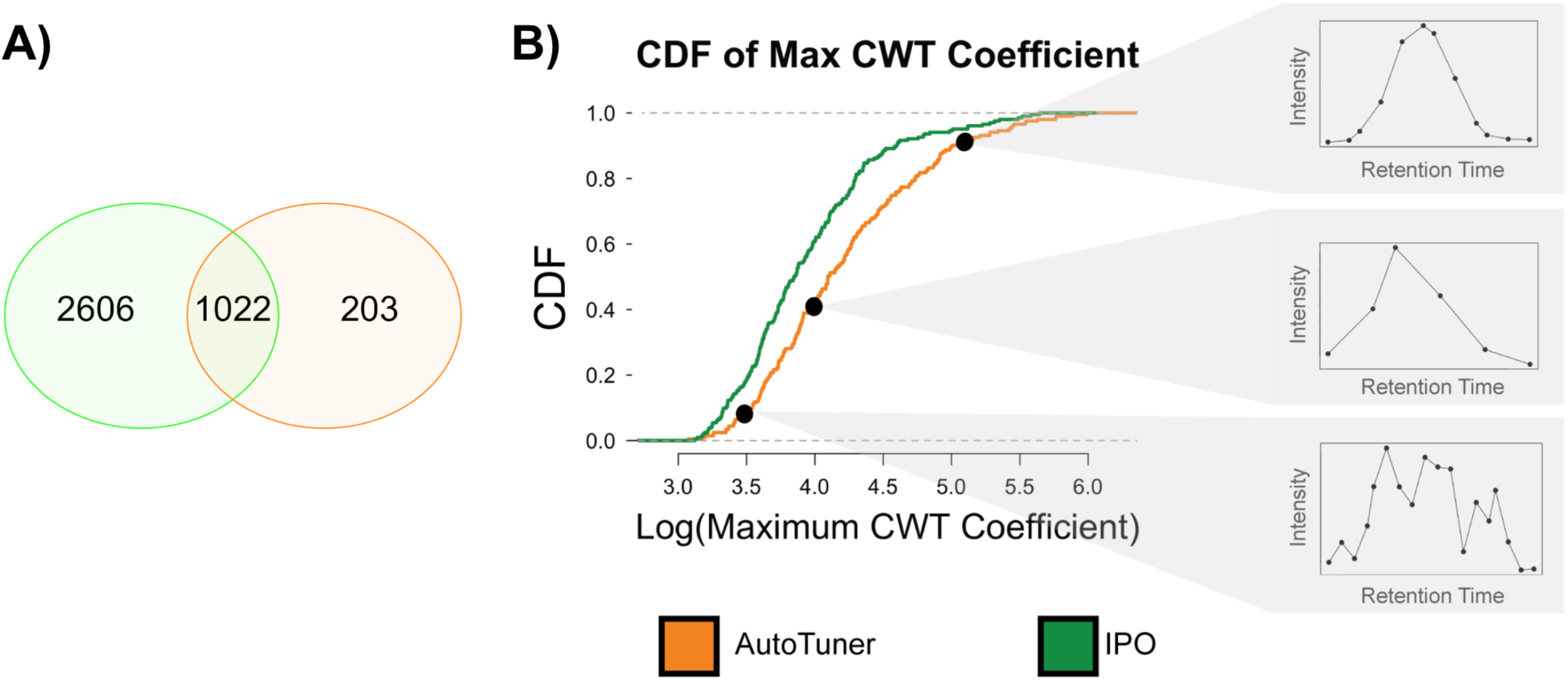
Comparing the differences between positive ion mode data generated by AutoTuner and IPO on culture dataset. A) portrays the overlap in the number of m/z-rt features generated by both methods. Features with an error of 5 ppm and retention time error of 20 seconds are placed in the intersect. B compares the differences in structural properties for the maximum continuous wavelet transform coefficient (CWT) between peaks detected only within AutoTuner (orange) and IPO (green). Both curves are empirical cumulative distribution functions (CDF) of the calculated metric. The inset shows three randomly selected EIC peaks that fall on distinct regions of the maximum CWT empirical cumulative distribution function to demonstrate how this metric influences peak quality. The curves were significantly different (KS-test, p < 10^−4^, n = 203). Results for positive ion mode data area CDF and negative ion mode data were similar and are found in figure S3 and S4, respectively.

We then compared the structural differences between features exclusively detected using IPO- and AutoTuner-derived parameters. We created an empirical cumulative distribution function (CDF) to compare the distribution of peak areas (Figure S3, Figure S4B) and maximum observed continuous wavelet transform (CWT) coefficients (Figure 2B, Figure S4C) of all EIC peaks belonging to features outside of the intersect. The maximum observed CWT coefficient increases with peak steepness, hence provides a measure of a peak’s chromatographic resolution (Figure 2: see inset). The CDF of each metric was significantly different in positive (KS-Test, Area: p < 10^−6^; CWT: p < 10^−4^, n = 203) and negative (KS-Test, area: p < 10^−14^, CWT: p < 10^−8^, n = 540) ionization mode data. Applying the same test on unbalanced comparisons (e.g., negative ion mode: 3420 IPO-vs. 540 AutoTuner-unique features) was more highly significant than using equivalent numbers of features obtained through subsampling.

Next, we used the abundance of MS^2^ spectra within each unique dataset to compare the abundance of real features (i.e., not noise) across feature tables. In total, there were more features with MS^2^ spectra from the feature table generated with IPO-derived parameters compared to AutoTuner-derived parameters (positive: 448 vs. 115; negative: 686 vs. 121, both for IPO vs. AutoTuner, respectively). However, when the features with MS^2^ spectra were normalized by the total number of features, AutoTuner-specific features contained more MS^2^ spectra than IPO-specific features (Figure 3). Unlike AutoTuner specific features, IPO specific features were significantly depleted for MS^2^ spectra relative to the intersect region (Hypergeometric Test, Negative ion mode: p < 10^−10^, Positive ion mode: p < 10^−10^).

**Figure 3.**
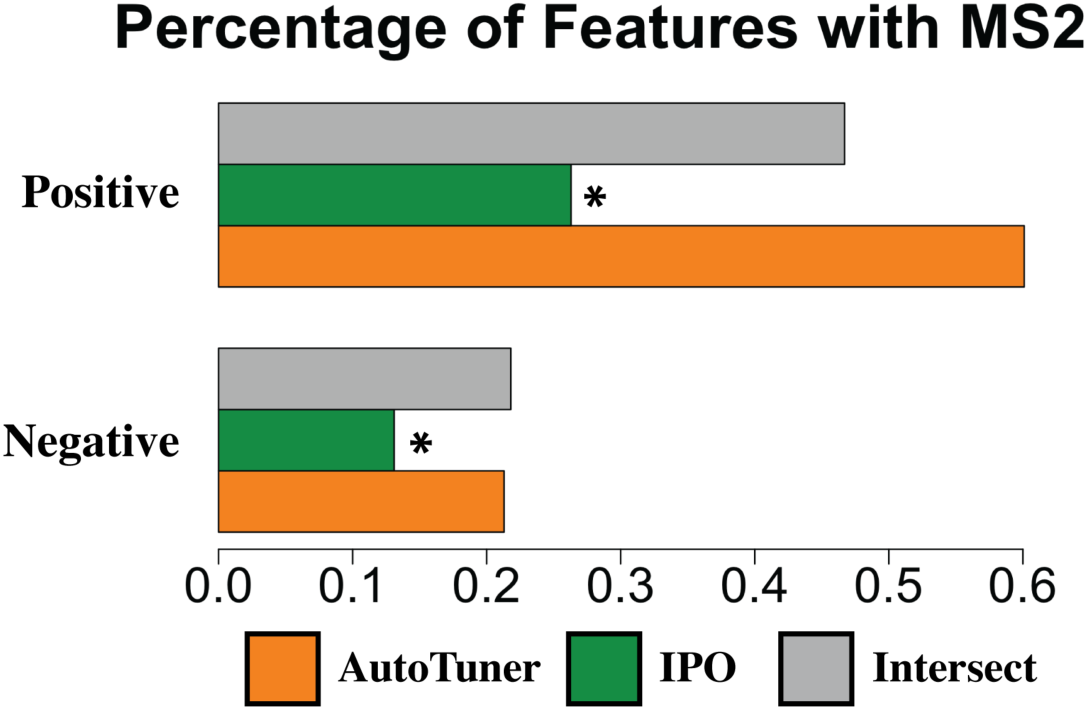
Percentage of MS^2^ spectra from feature tables generated by AutoTuner- and IPO-derived parameters. Each bar represents the percentage of features with MS^2^ spectra only detected by AutoTuner-derived parameters (orange); those detected only using IPO-derived parameters (green); or those detected within both feature tables (grey). (*) = groups significantly depleted in MS^2^ spectra relative to the intersect (Hypergeometric Test, p < 10^−10^).

Finally, using all three test datasets, we compared the time required to run AutoTuner and IPO (Table 3). After accounting for number of CPUs, AutoTuner ran hundreds to thousands of times faster.

**Table 3.** Run-times for AutoTuner and IPO required to run 6 common samples collected in positive (+) and negative (-) ionization modes. All system time measurements were done on an 8 CPUs and 10Gb of memory Ubuntu Xenial 16.04 Google Cloud instance. IPO ran on 8 CPUs, while AutoTuner ran on 1 CPU. The ratio accounts for the total computing power used to run both algorithms.

| Algorithm | Culture (-) | Culture (+) | Standards (-) | Standards (+) | Community (-) | Community (+) |
| --- | --- | --- | --- | --- | --- | --- |
| AutoTuner | 2 min | 9 min | 2 min | 3 min | 25 min | 26 min |
| IPO | 7 hr 23 min | 28 hr 40 min | 31 hr 56 min | 28 hr 5 min | 38 hr 4 min | 21 hr 27 min |
| Ratio (Auto/IPO) | 1479 | 1518 | 6970 | 4238 | 715 | 396 |

### Testing Robustness of AutoTuner Estimation

Figure 4 shows coefficients of variation and estimates for each 11-sample subset for the parameter ***ppm*** obtained from Monte Carlo analysis on culture and community datasets. Figures S5-S12 show the complete set of results from the Monte Carlo analysis for all parameters. For all parameter estimates, the variability of estimation decreased linearly with the number of samples used under both ionization modes (Figure 4A and Figures S5-S12). The rendered parameter estimates were consistent with expectations based on the mass analyzer used to generate the data (Table 2). With the exception of the ***Maximum Peak-width*** parameter estimate in the community dataset and the ***Noise*** estimate in the negative culture data, all parameters had a coefficient of variation (CV) less than or equal to 0.1 when using 9 samples to obtain estimates (Figures S5, S7, S9, and S11).

**Figure 4.**
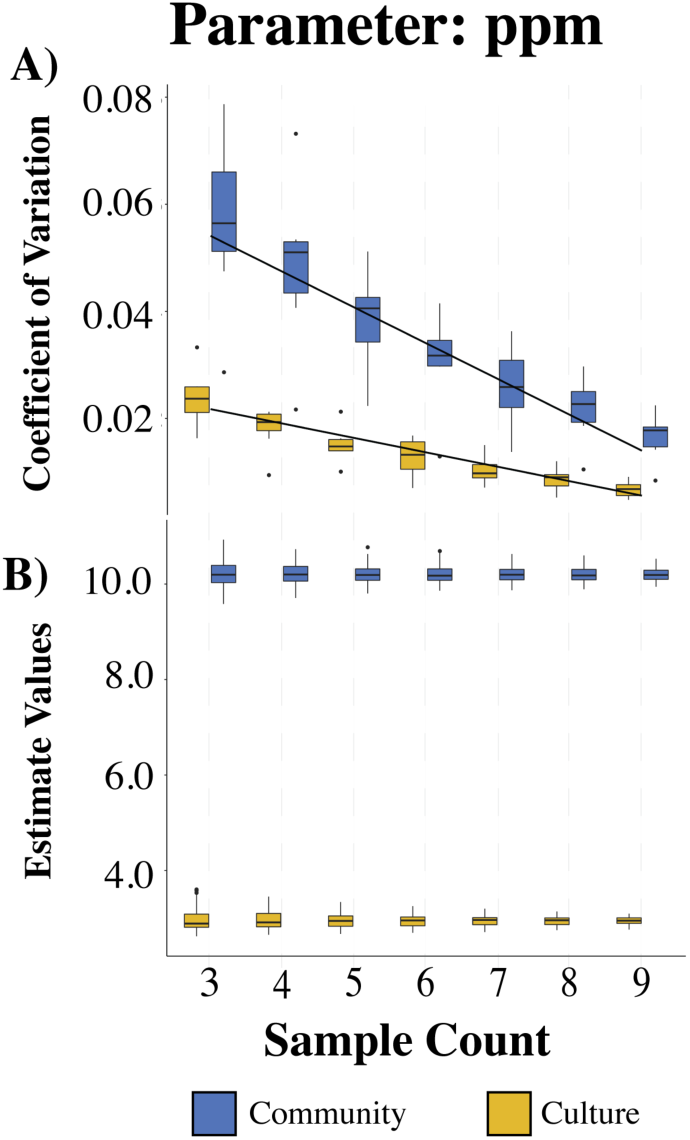
Results from Monte Carlo experiment for parameter ***ppm*** in positive ion mode data. A) depicts the distribution of coefficient of variation from parameters within 11 sample groups, while B) shows the distribution of all estimates for ***ppm***. Blue bars describe data collected by a qTOF instrument (community data) while yellow bars describe ultrahigh resolution (FT-ICR-MS) data (culture data). See figures S5-12 for other parameter estimates in each ionization mode.

## Discussion

AutoTuner is a robust, rapid, and high-fidelity estimator of untargeted mass spectrometry data processing parameters. Its unique design improves upon previous methods by providing a scalable framework to handle large datasets, reducing runtime, and generating high-accuracy parameters that retain known features. AutoTuner’s ease of use make it an ideal candidate to include within existing data processing pipelines.^39–42^ AutoTuner is available through BioConductor, and additional contributions are possible and encouraged.

AutoTuner’s high accuracy indicates that its parameter selection is based on true data features. One possible explanation for the lower accuracy of IPO is that the peak-width of the missing standards was below the ***Minimum Peak-width*** parameter selected by IPO (Table S2 and Figure S2).

AutoTuner parameter estimates were robust across all datasets and ionization modes. Some parameters like ***ppm, Noise, S/N Threshold, Prefilter intensity***, and ***Scan count*** reflect systematic properties inherent to the platform chromatography, mass analyzer, and/or sample matrix.^42^ Other parameters like ***Maximum peak-width*** are more specific to each sample; hence increasing the number of samples used to estimate parameters always strengthened their robustness. The low CV values for parameter estimates suggests that using a subset of samples to generate estimates returns a set representative for all samples. Based on our results, we recommend the use of 9 to 12 samples to generate estimates in culture and community datasets, respectively. For most of the parameters estimated here, 9 samples proved sufficient to obtain estimates with CV values less than 0.1. The 12-parameter recommendation originated from extrapolating the linear fits of these data to obtain 0.1 CV values for remaining parameters that failed this criterion (Figure S5, S7, S9, and S11). We were unable to check the robustness of the ***Group difference*** parameter estimate, as this parameter is estimated through a non-automatable cross sample comparison during the TIC peak detection step of the algorithm.

Although other algorithms return more parameter estimates than AutoTuner, the parameters calculated by AutoTuner represent continuous valued ones with the greatest possible number of choices. Performing a parameter sweeping optimization like previous approaches to estimate the remaining parameters reduces the total combinations of available centWave parameters from at least 2^4^* 5^8^ possible choices of parameters to 40. This is because the centWave algorithm, used by both XCMS and MZmine2 data processing tools, requires tuning of 11 distinct parameters. Of these, 8 are continuously valued, meaning that they can be any real number. The remainder are either boolean values or can be one of a few discrete choices (Table S4). The reduction of the total number of combinations is achieved by optimizing 7 of the 8 continuous valued parameters. In regards to the last continuous parameter not optimized by AutoTuner, mzDiff, we performed a sensitivity analysis to show that distinct values had minimal effect on the returned feature table (Table S5). Future contributions towards AutoTuner’s design can help the estimation of additional parameters.

AutoTuner’s low runtime indicates that the algorithm is scalable (Table 3). As more data is generated due to increases in analytical throughput and dataset size, AutoTuner will remain a tractable option to generate estimates of metabolomics data processing parameters.^4,44^ Because AutoTuner estimates parameters much faster than IPO, and IPO was shown to perform at a faster rate than software preceding it, we surmise that AutoTuner is the fastest parameter selection algorithm available.^22^

When considering the unique features identified by each algorithm in the culture dataset, AutoTuner found fewer features in each case (Figure 2, Figure S3). However, the uniquely detected AutoTuner features were enriched in MS^2^ spectra, an indicator of feature integrity. The significantly depleted number of MS^2^ spectra in features observed uniquely with IPO-derived parameters corroborates this claim. While identifying each distinct feature was beyond the scope of this study, these results suggest that AutoTuner-derived parameters generate smaller datasets with more information per feature. The size of the processed data using AutoTuner-derived parameters was in line with previous work validating the metabolome of *Escherichia coli* after performing stringent isotope labeling experiments and quality control filtering.^26^ The paucity of size-validated metabolome datasets precludes further evaluation of the feature number comparison.

AutoTuner has several avenues for possible improvement. First, AutoTuner could be parallelized to reduce computation time by a factor of the total number of CPUs used. Second, additional algorithms may be implemented to optimize parameters not covered here. The speed gained in the computation through AutoTuner’s “divide-and-conquer” approach comes at a loss of comparing EIC peaks across samples to estimate retention time correction algorithms. This challenge leaves room for the implementation of additional algorithms. Third, the replacement of the sliding window analysis with a more sophisticated and sensitive peak detection approach may eliminate the need for user input during the first portion of AutoTuner. However, this automation comes at the cost of manual inspection of the raw data. We support manual inspection of the raw data, because it provides a quality control check for the data generation steps leading up to the analysis. Despite these minor caveats, AutoTuner is a viable and time-saving option to determine proper data processing parameters for untargeted metabolomics data.

## Supporting information

Manuscript Supplementary Material

## Abbreviations

TIC: Total Ion Chromatogram.
EIC: Extracted Ion Current.
S/N: Signal to Noise.
KDE: Kernel Density Estimator.
KLD: Kullback-Leibler Divergence.
ppm: Parts Per Million.

## Acknowledgements

We thank Titus Brown and Ben Temperton for advice on algorithm validation, Arthur Eschenlauer for constructive feedback on software design, Krista Longnecker for continuous support and discussions, Gabriel Leventhal for mathematics advice and users of AutoTuner for debugging help through Github. Funding support included GEM and NSF graduate research fellowship program fellowships (CM) and grants from the MIT Microbiome Center (Award 6936800, EBK) and the Simons Foundation (Award ID #509034 EBK).

