## Supplementary material for "AutoTuner: High fidelity, robust, and rapid parameter selection for metabolomics data processing": Manuscript Supplementary Material

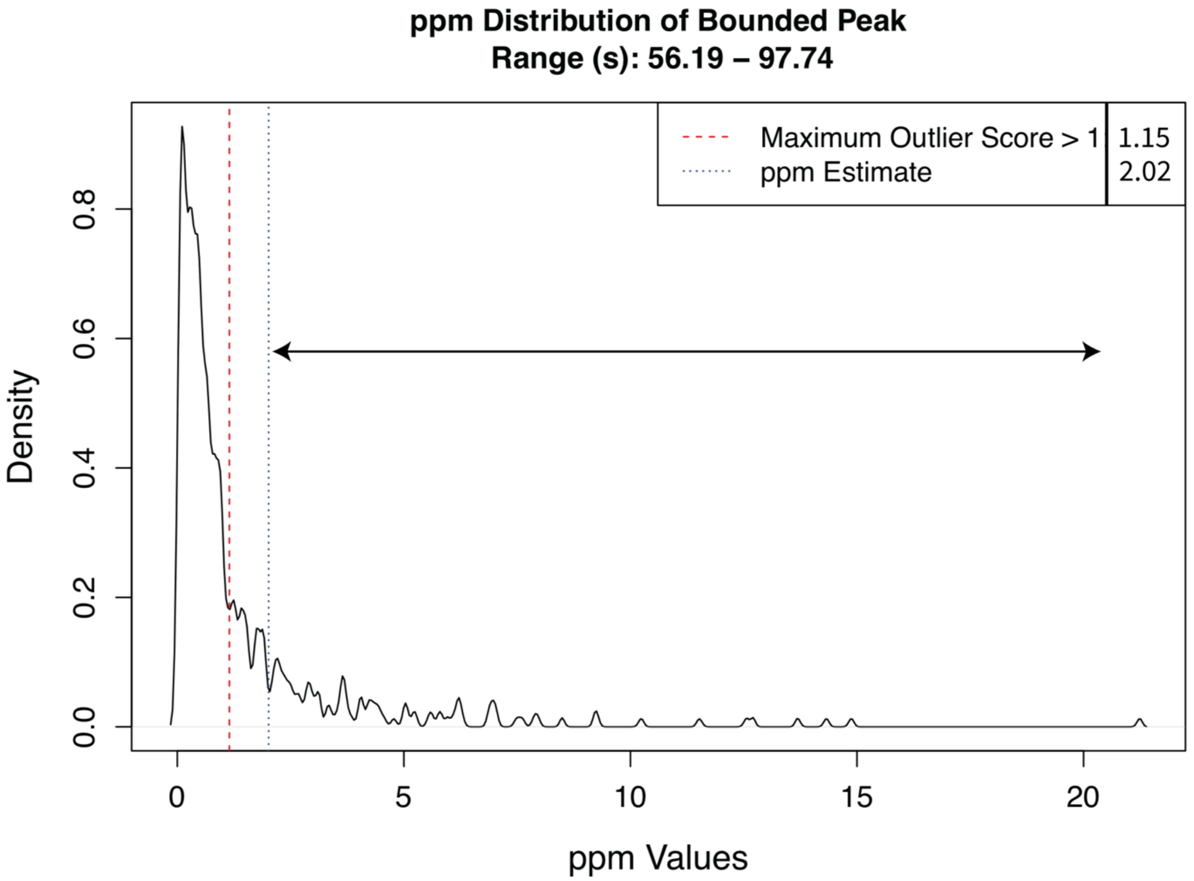


**Figure S1.** Example of AutoTuner-generated ppm error distribution. Such plots are returned by the algorithm to check quality of estimates. Red line represents the maximum ppm error value with an outlier score greater than 1 (see equation 3). In this example, a ppm error value of 1.15 meets this criterion (see legend). Blue line represents the ***ppm*** error parameter estimate described in equation 4, or 2.02 in this example (see legend). The "Range" subtitle represents the original chromatographic bounds of the TIC peak used to obtain estimates. The peaks under the arrow are assumed to originate from ppm values calculated from random associations of noise rather than from true features.

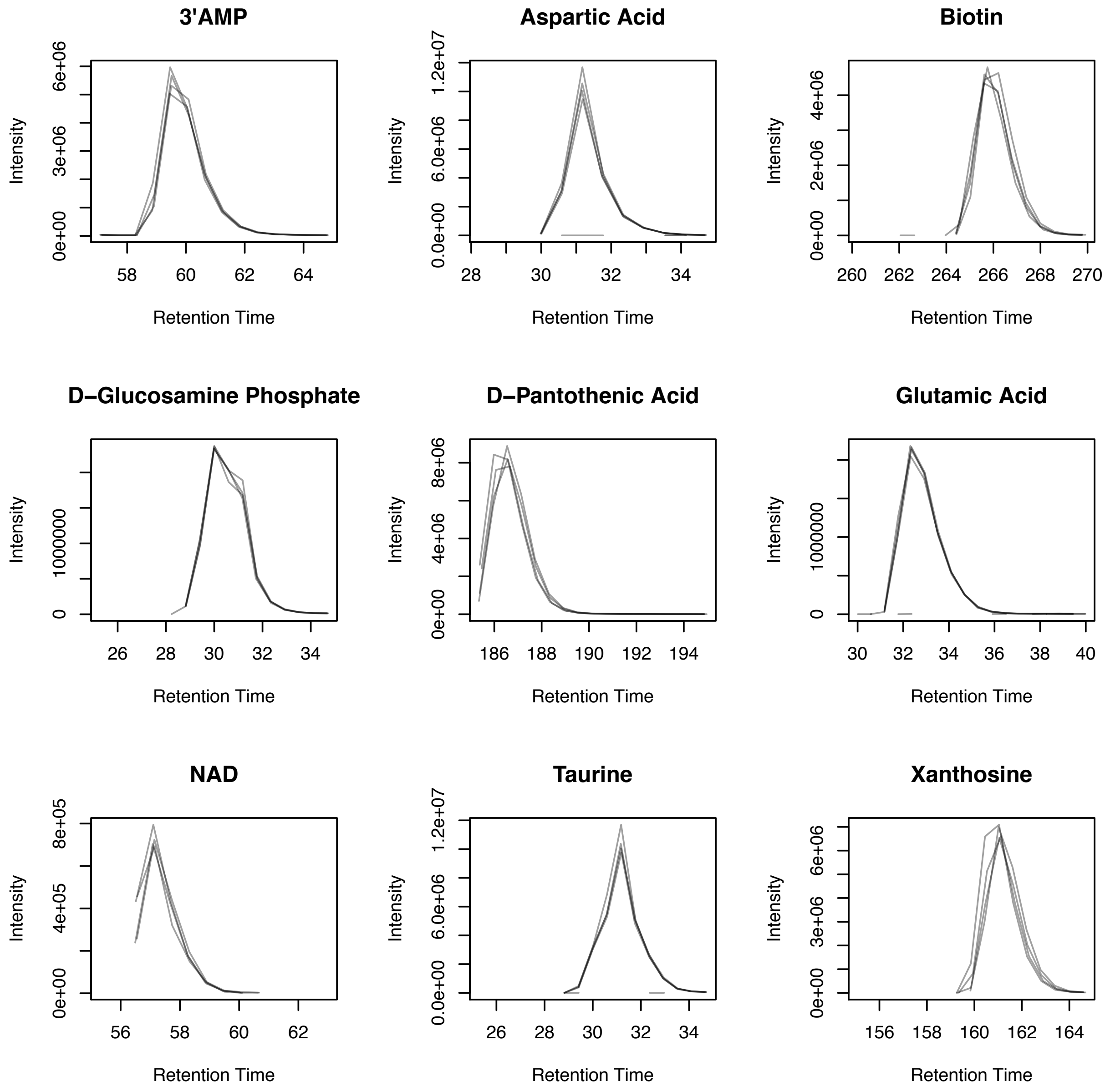


**Figure S2.** Example EIC peaks of standards not detected within feature table generated with IPO-derived parameters. The lines represent individual standard samples. 3’AMP = 3’-adenosine monophosphate; NAD = ß-nicotinamide adenine dinucleotide

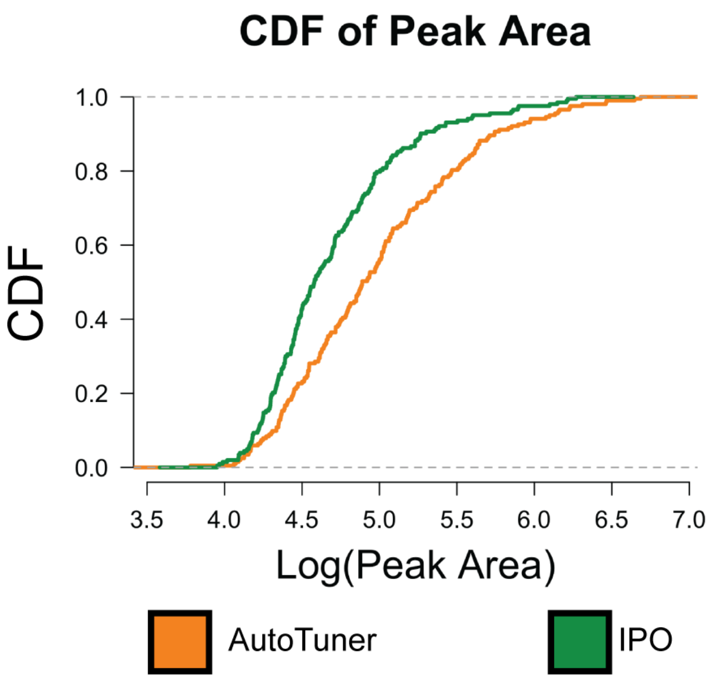


**Figure S3.** Positive ion mode data empirical cumulative distribution functions (CDF) comparison of peak area from EICs of features uniquely identified within feature tables generated with AutoTuner- and IPO-derived parameters. The curves were significantly different from one another (KS-test, Area: $p < 10^{-6}$; $n = 203$).

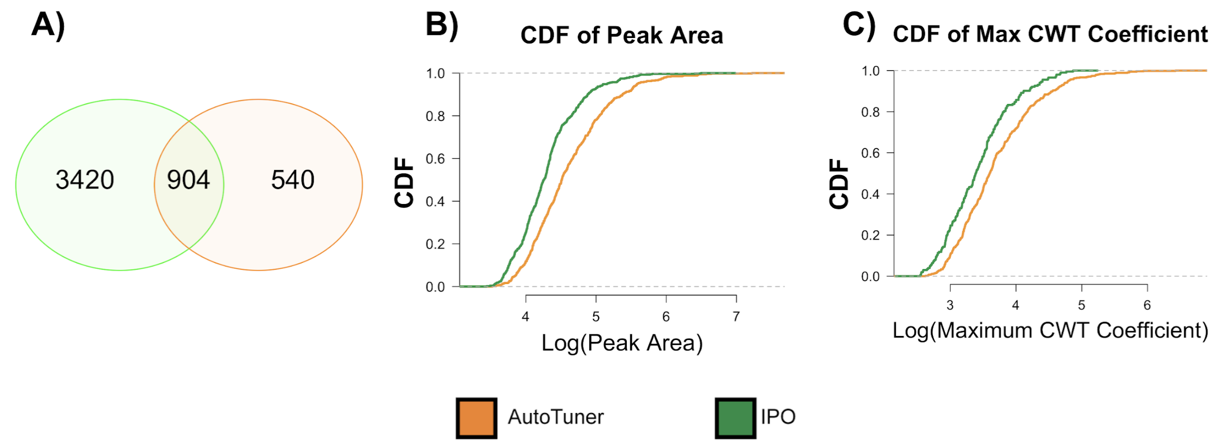


**Figure S4.** Negative ion mode data comparison of feature tables based on AutoTuner- and IPO-derived parameters on the culture dataset. A) portrays the overlap in the number of m/z-rt features generated by both methods. Features with an error of 5 ppm and retention time error of 20 seconds are placed in the intersect. B and C compare the differences in structural properties for the (B) peak area and (C) maximum continuous wavelet transform coefficient (CWT) between peaks detected only within AutoTuner or IPO. Both curves are empirical cumulative distribution functions (CDF) of the calculated metrics. An empirical cumulative distribution function is a non-parametric estimator of the underlying CDF of a random variable. In this case, the random variable is the set of calculated values for the AutoTuner- and IPO-specific features. CDFs for each metric were significantly different from one another (KS-test, Area: $p < 10^{-14}$; CWT: $p < 10^{-8}$, $n = 540$), similar to positive ion mode data.

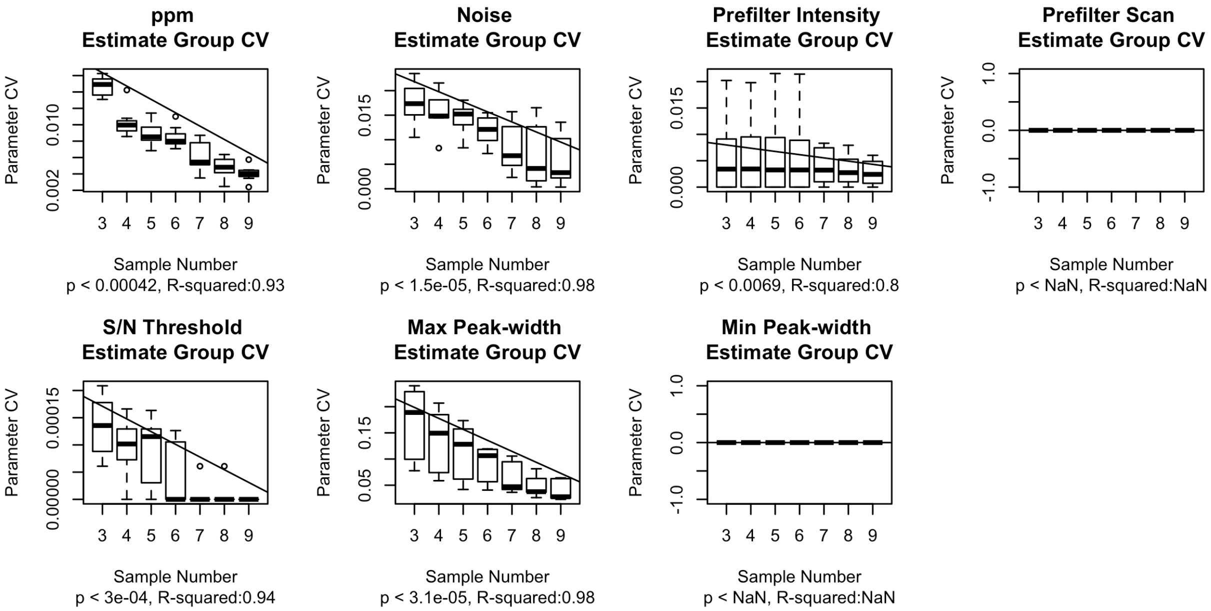


**Figure S5.** The coefficient of variation (CV) for groups of parameters estimated in the Monte Carlo analysis on negative mode community data. Each plot denotes the calculated CV values for each unique parameter. The x-axis describes the number of samples used to generate estimates, while the y-axis describes the CV of the estimates from each group of 11 randomly selected samples. P-value and $R^2$ statistics are derived from linear regressions of data ($n = 49$). (NaN = not a number).

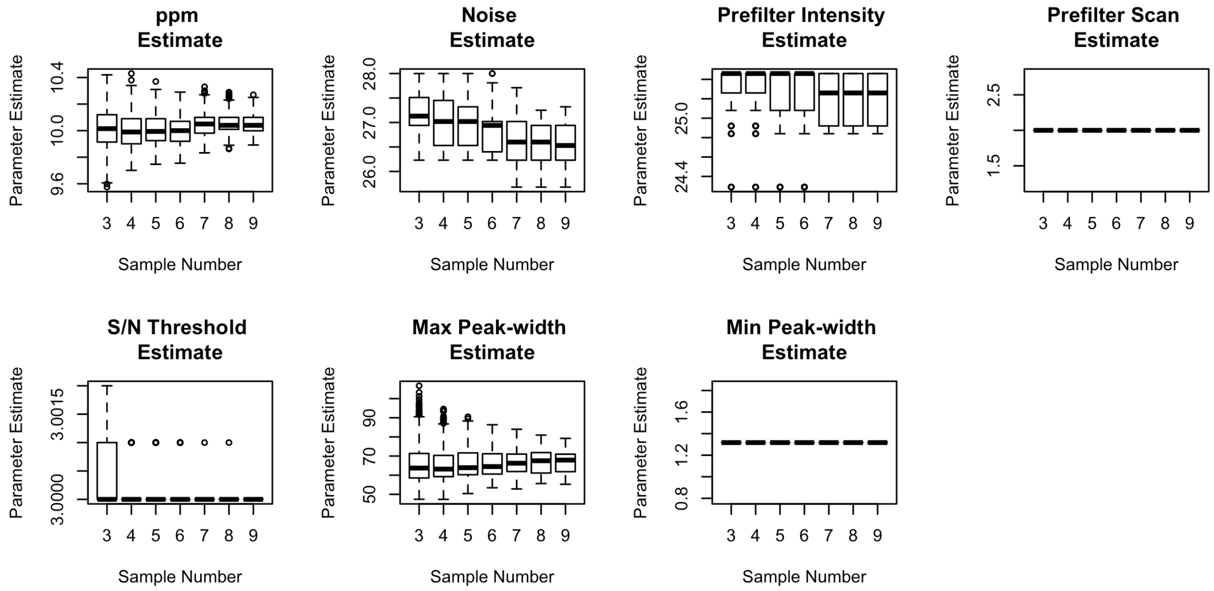


**Figure S6.** The parameters estimated in the Monte Carlo analysis on negative mode community data. Each plot denotes the calculated parameter estimate values for each unique parameter across 385 runs of AutoTuner. The x-axis describes the number of samples used to generate estimates, while the y-axis portrays the determined 55 parameter estimates within each n-sample subset (n = 3-9).

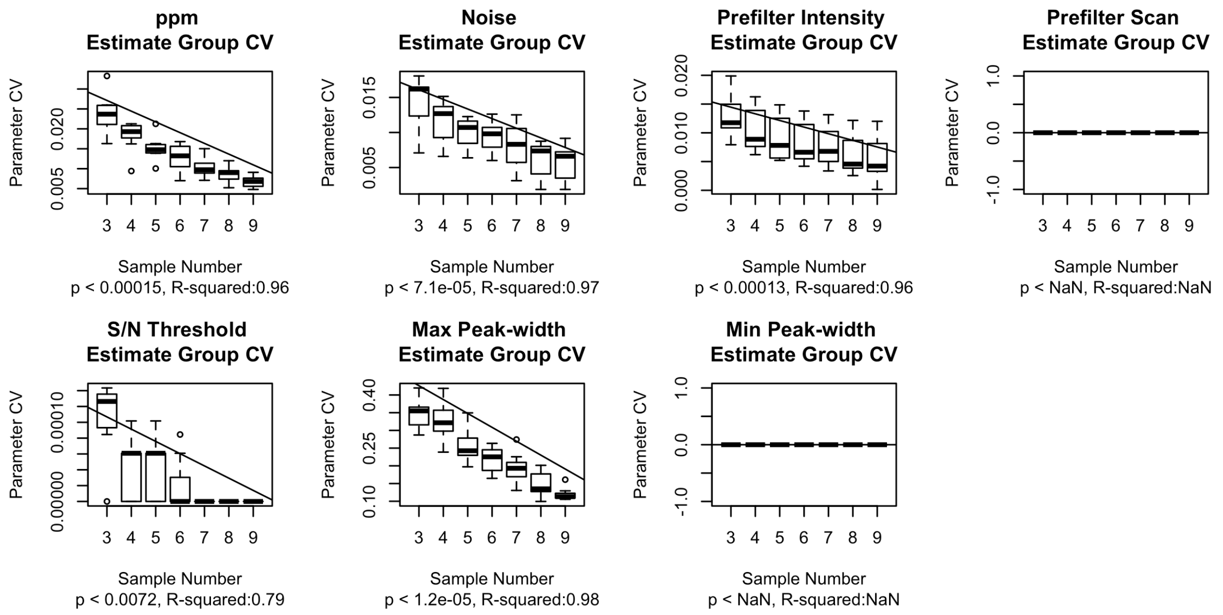


**Figure S7.** The coefficient of variation (CV) for groups of parameters estimated in the Monte Carlo analysis on positive mode community data. Each plot denotes the calculated CV values for each unique parameter. The x-axis describes the number of samples used to generate estimates, while the y-axis describes the CV of the estimates from each group of 11 randomly selected samples. P-value and $R^2$ statistics are derived from linear regressions of data (n = 49). (NaN = not a number).

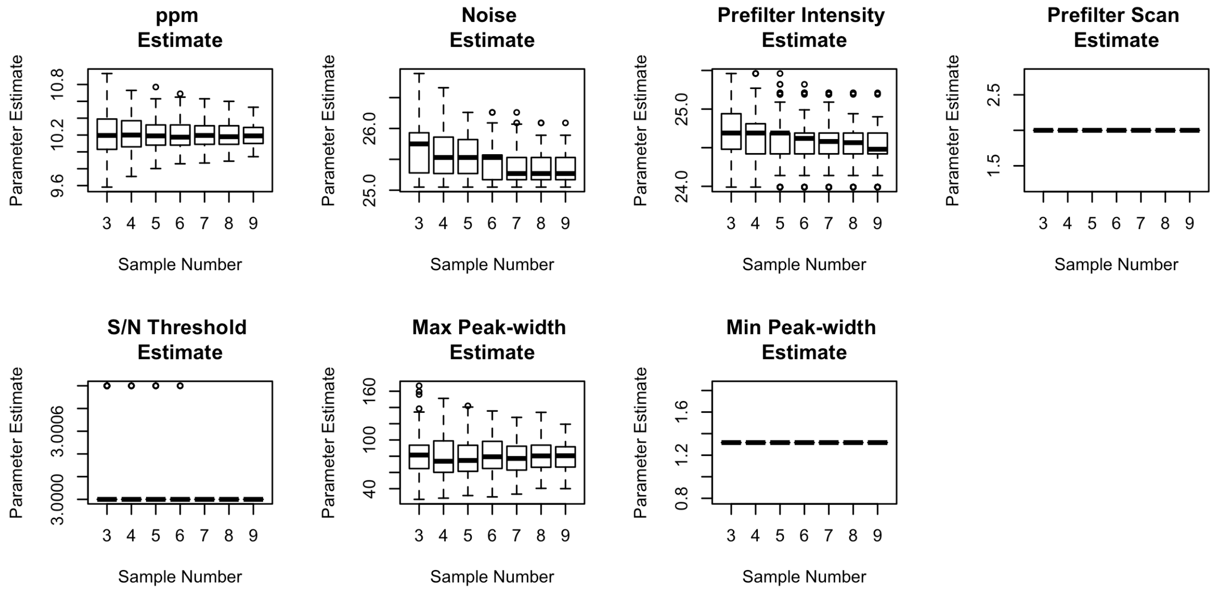


**Figure S8.** The parameters estimated in the Monte Carlo analysis on positive mode community data. Each plot denotes the calculated parameter estimate values for each unique parameter across 385 runs of AutoTuner. The x-axis describes the number of samples used to generate estimates, while the y-axis portrays the determined 55 parameter estimates within each n-sample subset (n = 3-9).

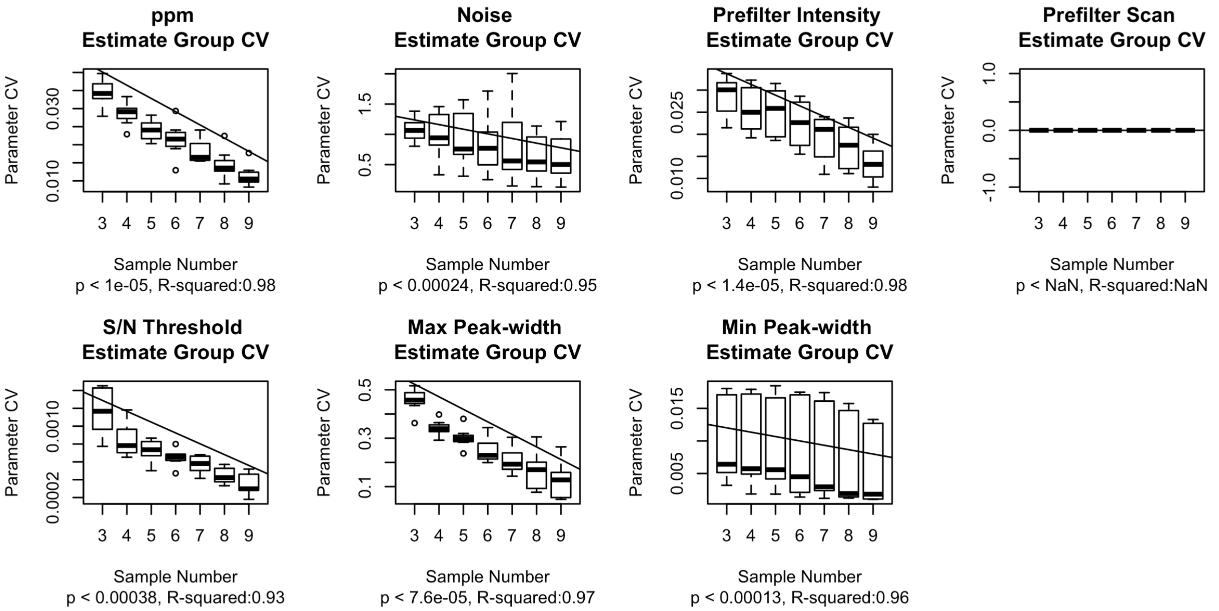


**Figure S9.** The coefficient of variation (CV) for groups of parameters estimated in the Monte Carlo analysis on negative mode culture data. Each plot denotes the calculated CV values for each unique parameter. The x-axis describes the number of samples used to generate estimates, while the y-axis describes the CV of the estimates from each group of 11 randomly selected samples. P-value and $R^2$ statistics are derived from linear regressions of data (n = 49). (NaN = not a number).

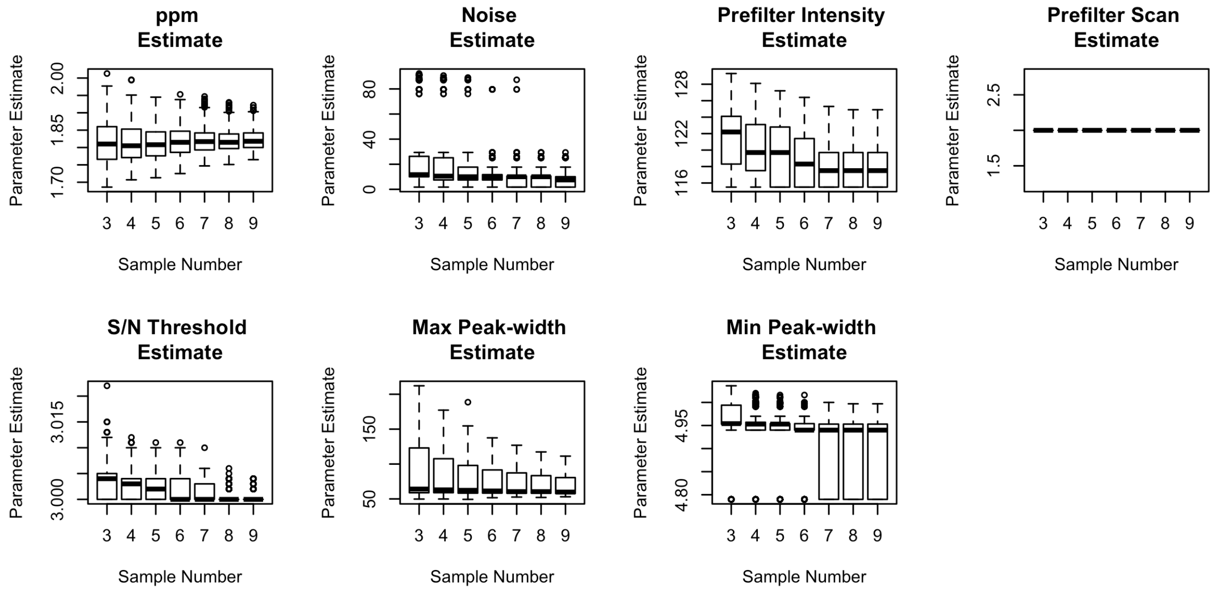


**Figure S10.** The parameters estimated in the Monte Carlo analysis on negative mode culture data. Each plot denotes the calculated parameter estimate values for each unique parameter across 385 runs of AutoTuner. The x-axis describes the number of samples used to generate estimates, while the y-axis portrays the determined 55 parameter estimates within each n-sample subset (n = 3-9).

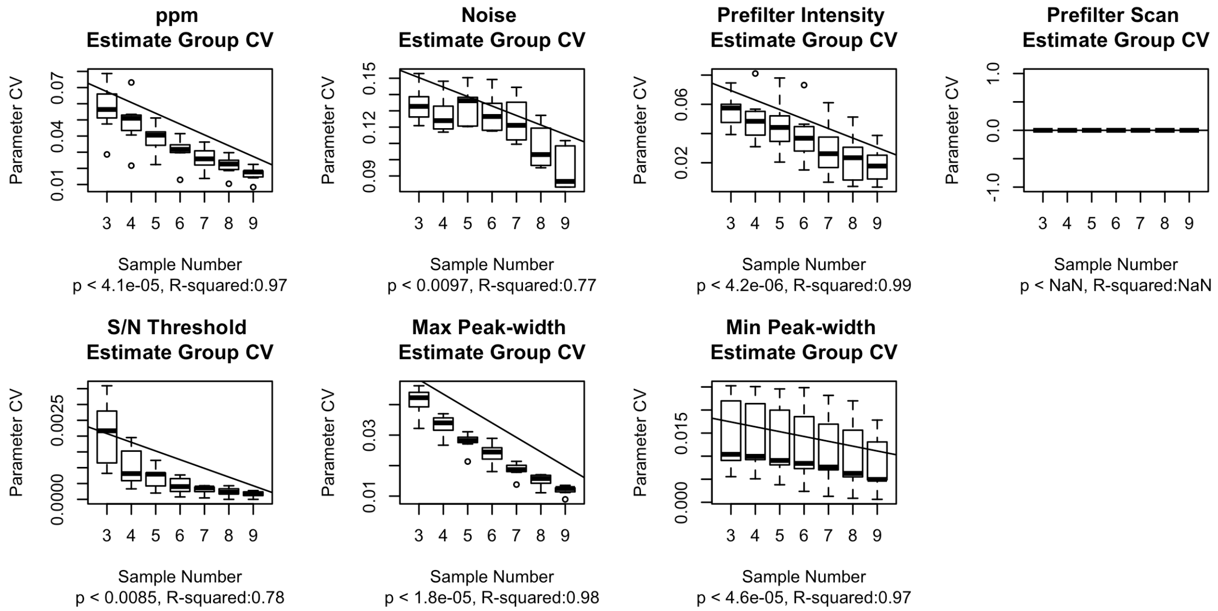


**Figure S11.** The coefficient of variation (CV) for groups of parameters estimated in the Monte Carlo analysis on positive mode culture data. Each plot denotes the calculated values for each unique parameter. The x-axis describes the number of samples used to generate estimates, while the y-axis describes the CV of the estimates from each group of 11 randomly selected samples. P-value and $R^2$ statistics are derived from linear regressions of data (n = 49). (NaN = not a number).

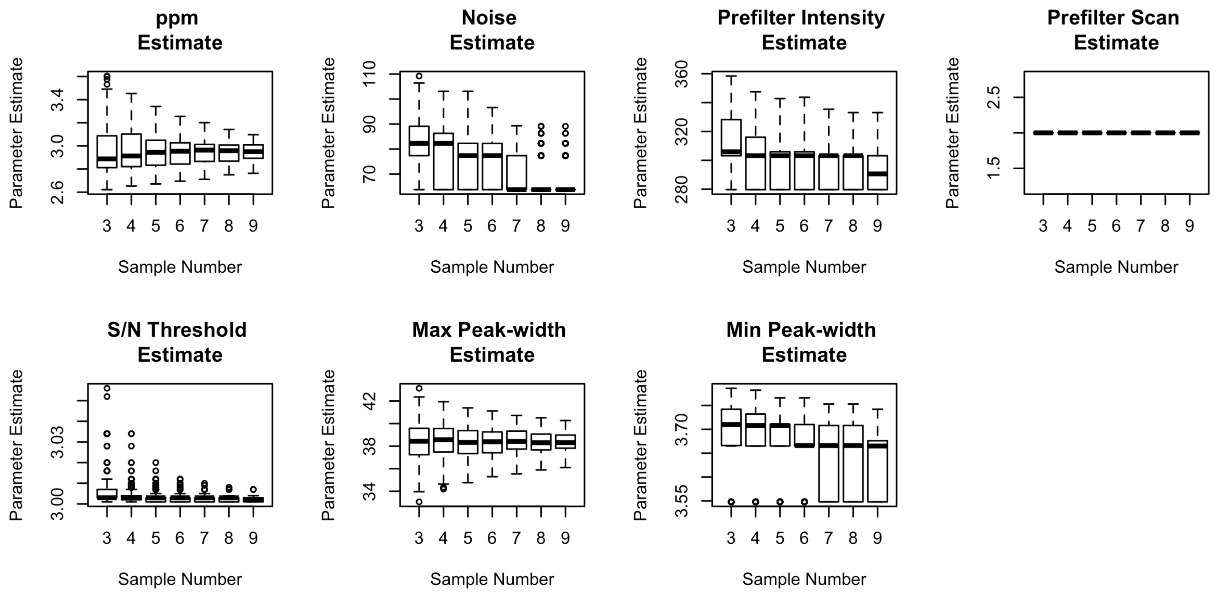


**Figure S12.** The parameters estimated in the Monte Carlo analysis on positive mode culture data. Each plot denotes the calculated parameter estimate values for each unique parameter across 385 runs of AutoTuner. The x-axis describes the number of samples used to generate estimates, while the y-axis portrays the determined 55 parameter estimates within each n-sample subset (n = 3-9).

**Table S1.** Standards used to validate AutoTuner accuracy. These compounds are common targets of metabolism and are commonly detected within untargeted metabolomics experiments. Compounds detected in both ionization modes are separated by "|" in the order they were presented in the "Ionization Mode" column.

| Compound | Ionization Mode | In AutoTuner | In IPO |
|---|---|---|---|
| 3-methyl-2-oxopentanoic acid | NEG | TRUE | TRUE |
| 3-methyl-2-oxobutanoic acid | NEG | TRUE | TRUE |
| 4-aminobenzoic acid | POS | TRUE | TRUE |
| 4-hydroxybenzoic acid | NEG | TRUE | TRUE |
| 4-methyl-2-oxopentanoic acid | NEG | TRUE | TRUE |
| adenosine 5'-monophosphate (5'AMP) | NEG\|POS | TRUE\|TRUE | FALSE\|TRUE |
| adenosine 3'-monophosphate (3'AMP) | NEG\|POS | TRUE\|TRUE | FALSE\|TRUE |
| 6-phosphogluconic acid | NEG | TRUE | TRUE |
| acetyl taurine | NEG | TRUE | TRUE |
| adenine | NEG\|POS | TRUE\|TRUE | TRUE\|TRUE |
| adenosine | POS | TRUE | TRUE |
| alpha-ketoglutaric acid | NEG | TRUE | TRUE |
| 4-amino-5-aminomethyl-2-methylpyrimidine (AmMP) | POS | TRUE | TRUE |
| arginine | POS | TRUE | FALSE |
| aspartic acid | NEG\|POS | TRUE\|TRUE | FALSE\|TRUE |
| biotin | NEG\|POS | TRUE\|TRUE | FALSE\|FALSE |
| caffeine | POS | TRUE | TRUE |
| citric acid | NEG | TRUE | TRUE |
| cytosine | POS | TRUE | TRUE |
| desthiobiotin | NEG\|POS | TRUE\|TRUE | TRUE\|TRUE |
| glucosamine phosphate | NEG | TRUE | FALSE |
| pantothenic acid | NEG\|POS | TRUE\|TRUE | FALSE\|TRUE |
| ribose 5-phosphate | NEG | TRUE | TRUE |
| 3-phosphoglyceric acid | NEG | TRUE | TRUE |

| | | | |
|---|---|---|---|
| diacetylchitobiose | POS | TRUE | TRUE |
| dihydroxy acetone phosphate | NEG | TRUE | TRUE |
| dimethylsulfoniopropionate (DMSP) | POS | TRUE | TRUE |
| ectoine | POS | TRUE | TRUE |
| folic acid | NEG\|POS | TRUE\|TRUE | TRUE\|TRUE |
| fosfomycin | NEG | TRUE | TRUE |
| fumarate | NEG | TRUE | TRUE |
| gamma-aminobutyric acid (GABA) | POS | TRUE | TRUE |
| glucose 6-phosphate | NEG | TRUE | TRUE |
| glutamic acid | NEG | TRUE | FALSE |
| glutamine | POS | TRUE | FALSE |
| glycine betaine | POS | TRUE | TRUE |
| glyphosate | NEG | TRUE | TRUE |
| guanine | POS | TRUE | TRUE |
| guanosine | NEG\|POS | TRUE\|TRUE | TRUE\|TRUE |
| 4-methyl-5-thiazoleethanol (HET) | POS | TRUE | TRUE |
| (4-amino-2-methyl-5-pyrimidinyl)methanol (HMP) | POS | TRUE | TRUE |
| indole 3-acetic acid | POS | TRUE | TRUE |
| inosine | NEG | TRUE | TRUE |
| inosine 5'-monophosphate | NEG\|POS | TRUE\|TRUE | TRUE\|TRUE |
| isethionic acid | NEG | TRUE | TRUE |
| citrulline | POS | TRUE | TRUE |
| glutathione | POS | TRUE | TRUE |
| glutathione oxidized | POS | TRUE | TRUE |
| isoleucine | POS | TRUE | TRUE |
| kynurenine | POS | TRUE | TRUE |
| leucine | POS | TRUE | TRUE |
| phenylalanine | POS | TRUE | TRUE |
| tryptophan | POS | TRUE | TRUE |
| tyrosine | POS | TRUE | TRUE |

| | | | |
|---|---|---|---|
| methionine | POS | TRUE | FALSE |
| 5'methylthioadenosine (MTA) | POS | TRUE | TRUE |
| muramic acid | NEG | TRUE | TRUE |
| N-acetyl d-glucosamine | POS | TRUE | TRUE |
| N-acetyl l-glutamic acid | NEG | TRUE | TRUE |
| N-acetylmuramic acid | NEG\|POS | TRUE\|TRUE | TRUE\|TRUE |
| ß-nicotinamide adenine dinucleotide (NAD) | NEG\|POS | TRUE\|TRUE | FALSE\|FALSE |
| ß-nicotinamide adenine dinucleotide phosphate (NADP) | NEG | TRUE | TRUE |
| ornithine | POS | TRUE | TRUE |
| orotic acid | NEG | TRUE | TRUE |
| phosphoenolpyruvate | NEG | TRUE | TRUE |
| proline | POS | TRUE | TRUE |
| pyridoxine | POS | TRUE | TRUE |
| riboflavin | POS | TRUE | FALSE |
| S-(1,2-dicarboxyethyl)glutathione | POS | TRUE | TRUE |
| S-(5'-adenosyl) -L-homocysteine (SAH) | NEG\|POS | TRUE\|TRUE | TRUE\|TRUE |
| S-adenosyl-l-methionine (SAM) | POS | TRUE | FALSE |
| serine | POS | TRUE | FALSE |
| sn-glycerol 3-phosphate | NEG\|POS | TRUE\|TRUE | TRUE\|TRUE |
| succinic acid | NEG | TRUE | TRUE |
| syringic acid | NEG | TRUE | TRUE |
| taurine | NEG | TRUE | FALSE |
| thiamine monophosphate | POS | FALSE | FALSE |
| threonine | POS | TRUE | TRUE |
| thymidine | NEG | TRUE | TRUE |
| triacetylchitotriose | POS | TRUE | TRUE |
| uracil | POS | TRUE | TRUE |
| uridine 5'-monophosphate | POS | TRUE | TRUE |

| | | | |
|---|---|---|---|
| valine | POS | TRUE | TRUE |
| xanthine | NEG\|POS | TRUE\|TRUE | TRUE\|TRUE |
| xanthosine | NEG\|POS | TRUE\|TRUE | FALSE\|TRUE |

**Table S2.** Parameters used to process data. We rounded the values returned by AutoTuner and IPO at the tenths place. Each column aside from the "Dataset" and "Method" represent XCMS parameters described in Table 1. The community dataset is not mentioned here, as no comparison between IPO- and AutoTuner-parametrized feature tables was performed. The same standard set of parameters were used for density grouping and loess spline retention time correction. XCMS function syntax is described in parentheses. For the first run of density grouping (group.density): group difference =10, minfrac = 0, minsamp = 1, mzwid = 0.001. For the second run of density grouping after retention time correction (group.density):, group difference = 5, minfrac = 0.5, minsamp = 1, mzwid = 0.001. For loess spline retention time correction (retcor.peakgroups): span = 0.5.

| **Dataset** | **Method** | ***Maximum Peak-width*** | ***Minimum Peak-width*** | ***ppm*** | ***Noise*** | ***Prefilter Intensity*** | ***Scan Count*** | ***S/N Threshold*** |
|---|---|---|---|---|---|---|---|---|
| Pos Standards | IPO | 26.0 | 12.0 | 6.2 | 250.0 | 100.0 | 3.6 | 10 |
| Pos Standards | AutoTuner | 29.3 | 5.7 | 4.0 | 436.8 | 1421.3 | 2.0 | 6 |
| Pos Culture | IPO | 48.0 | 18.6 | 5.3 | 470 | 100.0 | 2.5 | 7 |
| Pos Culture | AutoTuner | 38.3 | 3.6 | 3.0 | 66.7 | 292.0 | 2.0 | 3 |
| Neg Standards | IPO | 26.0 | 12.0 | 6.2 | 250.0 | 100.0 | 3.6 | 10 |
| Neg Standards | AutoTuner | 29.3 | 5.7 | 4.0 | 436.8 | 1421.3 | 2.0 | 6 |
| Neg Culture | IPO | 60.0 | 27.4 | 4.7 | 121.0 | 100.0 | 4.0 | 9 |
| Neg Culture | AutoTuner | 66.9 | 4.9 | 1.8 | 7.8 | 117.5 | 2.0 | 3 |

**Table S3.** Feature count from each dataset during the different stages of quality assurance processing of culture data. The initial feature count was reduced after processing to remove blanks ('post blank)', features found in only one replicate ('post reproducibility), isotopologues and adducts ('post isotopes', and 'post adducts', respectively), and features with a CV greater than 0.4 in the pooled samples ('post CV').

| **Ionization Mode** | **Algorithm** | **Initial Feature Count** | **Post Blank** | **Post Reproducibility** | **Post Isotopes** | **Post Adducts** | **Post CV** |
|---|---|---|---|---|---|---|---|
| Negative | IPO | 40422 | 37903 | 8225 | 7695 | 4324 | 4226 |
| Negative | AutoTuner | 22599 | 17640 | 2921 | 2805 | 1444 | 1363 |
| Positive | IPO | 28794 | 28042 | 5907 | 5591 | 3628 | 3520 |
| Positive | AutoTuner | 13731 | 12451 | 2099 | 2012 | 1225 | 1143 |

**Table S4.** Standard parameters used within centWave algorithm and their number of possible combinations. We cite these values in our discussion of speed improvements gained via AutoTuner relative to traditional parameter sweeping approaches dependent on optimization functions.

| Parameter | Type | Possible Choices | Checked by AutoTuner |
|---|---|---|---|
| ***ppm*** | Continuous | Infinite | Yes |
| ***S/N Threshold*** | Continuous | Infinite | Yes |
| ***Scan count*** | Continuous | Infinite | Yes |
| ***Noise*** | Continuous | Infinite | Yes |
| ***Prefilter intensity*** | Continuous | Infinite | Yes |
| ***Minimum Peak-width*** | Continuous | Infinite | Yes |
| ***Maximum Peak-width*** | Continuous | Infinite | Yes |
| ***mzDiff*** | Continuous | Infinite | No |
| ***Fit gauss*** | Boolean | 2 | No |
| ***Mz center function*** | Discrete | 4 | No |
| ***Integrate*** | Discrete | 2 | No |

**Table S5.** Number of unique features observed after processing data with unique mzDiff values. Columns two and three denote the mzDiff values used during pairwise comparisons of feature tables. Missing Count column represents the number of features observed outside the intersect of both feature tables. Feature tables were generated from 8 negative ion mode community data samples.

| Missing Count | mzDiff value of First Feature Table | mzDiff value of Second Feature Table |
|---|---|---|
| 0 | -0.001 | -0.002 |
| 0 | -0.002 | -0.003 |
| 0 | -0.003 | -0.004 |
| 0 | -0.004 | -0.005 |
| 0 | -0.005 | -0.006 |
| 0 | -0.006 | -0.007 |
| 0 | -0.007 | -0.008 |
